# The E3 ubiquitin ligase RNF216/TRIAD3 is a central regulator of the hypothalamic-pituitary-gonadal axis

**DOI:** 10.1101/2021.03.21.436306

**Authors:** Arlene J. George, Yarely C. Hoffiz, Christopher Ware, Bin Dong, Ning Fang, Erik Hrabovszky, Angela M. Mabb

## Abstract

RNF216/TRIAD3 is an E3 ligase that ubiquitinates substrates in the nervous system. Recessive mutations in *RNF216/TRIAD3* cause Gordon Holmes syndrome (GHS), where hypogonadotropic hypogonadism is a core phenotype. However, the functions of RNF216/TRIAD3 within the neuroendocrine system are not well-understood. Here, we used the CRISPR-Cas9 system to knock out *Rnf216/Triad3* in GT1-7 cells, a GnRH immortalized cell line derived from mouse hypothalamus. *Rnf216/Triad3* knockout cells had decreased steady state *Gnrh* and reduced calcium transient frequency. To address functions of RNF216/TRIAD3 *in vivo*, we generated a *Rnf216/Triad3* constitutive knockout (KO) mouse. KO mice of both sexes showed reductions in GnRH and soma size. Furthermore, KO mice exhibited sex-specific phenotypes with males showing gonadal impairment and derangements in gonadotropin release compared to KO females, which only had irregular estrous cyclicity. Our work shows that dysfunction of RNF216/TRIAD3 affects the HPG axis in a sex-dependent manner, implicating sex-specific therapeutic interventions for GHS.

**Highlights:**

- *Rnf216/Triad3* controls *Gnrh* and intrinsic hypothalamic cell activity
- *Rnf216/Triad3* knockout male mice have greater reproductive impairments than females
- *Rnf216/Triad3* controls the HPG axis at multiple levels

## Introduction

The integrity of the hypothalamic-pituitary-gonadal (HPG) axis is necessary for neuroendocrine control of reproductive behavior that systemically allows for the secretion of hormones via tightly regulated neural networks (Harris, 1955). Activation of the HPG axis is initiated by a population of kisspeptin neurons in the anteroventral periventricular nucleus (AVPV) and arcuate nucleus (ARN) that mediates the release of kisspeptin, which binds to G-protein coupled receptor 54 (GPR54) receptors located on the surface of gonadotropin-releasing hormone (GnRH) positive neurons in the preoptic area of the hypothalamus (Han et al., 2005; Herbison, 2016). The activation of these receptors facilitates calcium-dependent pathways that are critical for GnRH production and release (Armstrong et al., 2009; Kotani et al., 2001; Moenter et al., 2003). GnRH then stimulates secretory gonadotropes located in the anterior pituitary to release the gonadotropins, luteinizing hormone (LH) and follicle-stimulating hormone (FSH) to the gonads, which regulates the secretion of sex steroids (Plant, 2015).

Loss-of-function mutations in HPG axis neuropeptides, gonadotropins, and receptor genes cause hypogonadotropic hypogonadism (HH) (Achrekar et al., 2010; Bramble et al., 2016; Bruysters et al., 2008; de Roux et al., 2003; de Roux et al., 1997; Seminara et al., 2003). HH is a condition that is defined by gonadal impairments with decreased levels of sex steroids due to HPG axis defects (de Roux et al., 1997; Kalantaridou and Chrousos, 2002). HH is prevalent in a diverse set of disorders that include Kallmann syndrome, Prader-Willi syndrome, Fragile X syndrome, cerebellar ataxias and central and peripheral hypomyelination (Alsemari, 2013; Angulo et al., 2015; Hardelin, 2001; Timmons et al., 2006). One neurological disease with a defining feature of HH is Gordon Holmes syndrome (GHS), a rare disorder with a constellation of signs and symptoms that also include cerebellar ataxia, dysarthria, cognitive impairment, and neurodegeneration (Holmes, 1908; Seminara et al., 2002). GHS individuals are diagnosed around pubertal-age and often present with poor development of secondary sexual characteristics concurrent with low levels of LH and testosterone (Alqwaifly and Bohlega, 2016; Ganos et al., 2015; Mehmood et al., 2017; Quinton et al., 1999; Seminara et al., 2002). Clinical studies show that some, but not all GHS patients will respond to treatment with exogenous GnRH, but this remains a short-term solution as patient responsiveness wanes over time (Margolin et al., 2013; Quinton et al., 1999; Seminara et al., 2002). Clinical studies have revealed that some female patients experience a less severe HH phenotype than males with only one case of a woman having two successful pregnancies (Lieto et al., 2019). These findings indicate that there could be sex differences in the pervasiveness of HH in GHS.

Protein ubiquitination is an ATP-driven posttranslational modification that involves the covalent addition of a small protein called ubiquitin to other proteins resulting in a multitude of cellular functions (Hershko and Ciechanover, 1998). This is initiated by an E1 ubiquitin activating enzyme that transfers the ubiquitin to an E2 ubiquitin conjugating enzyme, which will then form a complex with an E3 ubiquitin ligase to transfer ubiquitin to the target protein or substrate (Komander and Rape, 2012; Zheng and Shabek, 2017). Emerging evidence indicate that protein ubiquitination is critical for HPG axis function. For example, heterozygous mutations in the E3 ligase, makorin RING-finger protein 3 (*MKRN3*), and low serum levels of MKRN3 are linked to central precocious puberty (Aycan et al., 2018; de Vries et al., 2014; Grandone et al., 2018; Liu et al., 2017; Stecchini et al., 2016). Similarly, disruptions in protein ubiquitin enzymes are associated with HH where homozygous recessive or compound heterozygous mutations in the E3 ligases *STUB1/CHIP* or *RNF216/TRIAD3* are causative for GHS (Alqwaifly and Bohlega, 2016; Calandra et al., 2019; Ganos et al., 2015; George et al., 2018; Hayer et al., 2017; Margolin et al., 2013; Mehmood et al., 2017; Santens et al., 2015; Sawyer et al., 2014; Song et al., 2013).

RNF216/TRIAD3 is a RING (Really Interesting New Gene) E3 ligase that encodes for multiple E3 ligase isoforms including TRIAD3A, TRIAD3B, TRIAD3C, and TRIAD3D/E (Chuang and Ulevitch, 2004). Notably, TRIAD3A can assemble lysine-48 (-K48) ubiquitin linkages leading to proteasome-dependent degradation of multifarious substrates including those involved in immunological function and cell death (Alturki et al., 2018; Chuang and Ulevitch, 2004; Fearns et al., 2006; Nakhaei et al., 2009). RNF216/TRIAD3 also ubiquitinates substrates that regulate autophagy (Kim et al., 2018; Wang et al., 2016; Xu et al., 2014). On the contrary, recent literature highlights a role for TRIAD3B in the assembly of -K63 ubiquitin linkages, which mediate signal transduction processes (Schwintzer et al., 2019; Seenivasan et al., 2019). All of these studies have been focused outside of the nervous system. The only documented role of RNF216/TRIAD3 in the nervous system is its regulation in learning-related synaptic plasticity. TRIAD3A localizes to clathrin-coated pits and is associated with endocytic zones within the postsynaptic membrane of primary neurons. Triad3A ubiquitinates ARC, an immediate early gene product, which leads to a reduction in surface α-amino-3-hydroxy-5-methyl-4-isoxazolepropionic acid (AMPA) receptors (Mabb et al., 2014). A loss in Triad3A-dependent ARC ubiquitination increases AMPA receptor endocytosis and prolongs metabotropic glutamate receptor-mediated long term depression (mGluR-LTD) in hippocampal neurons (Wall et al., 2018). Remarkably, there is no literature describing a role for *RNF216/TRIAD3* in HH, one of the core phenotypes associated with GHS, nor are there animal models that have yet attempted to dissect the functions of ubiquitin enzymes within an intact HPG axis.

In previous experiments, we found that Triad3A regulates the general trafficking of receptors in neurons. In this study, we unveiled a role of RNF216/TRIAD3 within the complex HPG axis. Moreover, RNF216/TRIAD3 was found to be expressed in the hypothalamus and in GT1-7 cells, an immortalized cell line derived from mouse GnRH neurons, which secretes GnRH (Mellon et al., 1990). We first developed an *in vitro* hypothalamic cellular model by knocking out all isoforms of *Rnf216/Triad3* from GT1-7 cells using the CRISPR-Cas9 system. We found that a loss of *Rnf216/Triad3* reduced *GnRH* production without changing the expression or localization of key GnRH neuron receptors. Using a genetically encoded calcium sensor, *Rnf216/Triad3* knockout (KO) cells exhibited reduced activity. Characterization of *Rnf216/Triad3* KO mice exhibited deteriorations in reproductive health that included lower breeding viability in males, irregular cyclicity of the female estrous cycle, and decreased male gonadal volume. Surprisingly, the downstream effector of GnRH, FSH was elevated in male but unchanged in female KO mice despite decreased GnRH and GnRH neuron size in both sexes. These findings indicate that the loss of function of *RNF216/TRIAD3* is associated with an impaired neuroendocrine axis and shows that dysfunction of RNF216/TRIAD3 *in vivo* affects HPG axis profiles in a sex-dependent manner.

## Results

### *Rnf216/Triad3* KO GT1-7 cells have decreased *GnRH* and calcium transient frequency

Since HH in GHS is manifested by decreased GnRH release (Alqwaifly and Bohlega, 2016; Ganos et al., 2015; Holmes, 1908; Mehmood et al., 2017; Quinton et al., 1999; Seminara et al., 2002), and the presence of feedback loops within the HPG axis can alter GnRH release (Plant, 2015), we took advantage of an isolated immortalized mature mouse hypothalamic cell line (GT1-7) that produces and secretes GnRH (Mellon et al., 1990) and utilized the CRISPR-Cas9-mediated knockout strategy (Shalem et al., 2014). We generated two separate gRNA clones (A and B) targeting all isoforms of the *Rnf216/Triad3* gene. Clone A decreased RNF216/TRIAD3 by 63 ± 0.10% whereas clone B yielded a 90 ± 0.03% decrease (*p*<0.0001, One-way ANOVA; Figure 1A). Purification of genomic DNA followed by Sanger sequencing of CRISPR Control, A, and B clones demonstrated gRNA target specificity within the *Rnf216/Triad3* targeted genomic regions (Figure 1B and Supplementary Figure 1A). Spectral decomposition using the Tracking of Indels by Decomposition (TIDE) tool (Brinkman et al., 2014), showed the successful creation of indels ranging from 1 to 10 base pairs for deletions and insertions (Supplementary Figure S1B). While CRISPR B lacked the presence of any wildtype (WT) *Rnf216/Triad3* sequence when compared to CRISPR control cells, there was still the presence of a small fraction of WT sequence (3.1%) in CRISPR A cells (Supplementary Figure S1B), which agreed with our ability to detect residual full-length RNF216/TRIAD3 protein in CRISPR A (Figure 1A). When measuring *Gnrh* using quantitative real-time polymerase chain reaction (qPCR), we found a graded decrease in *Gnrh* between control, CRISPR A, and CRISPR B that mirrored the changes in RNF216/TRIAD3 expression. Particularly, there was a significant decrease of *Gnrh* in CRISPR B cells (*p*=0.0342, One-way ANOVA; Figure 1C). These findings suggest that RNF216/TRIAD3 controls the production of *GnRH* in a dose-dependent manner.

**Figure 1:**
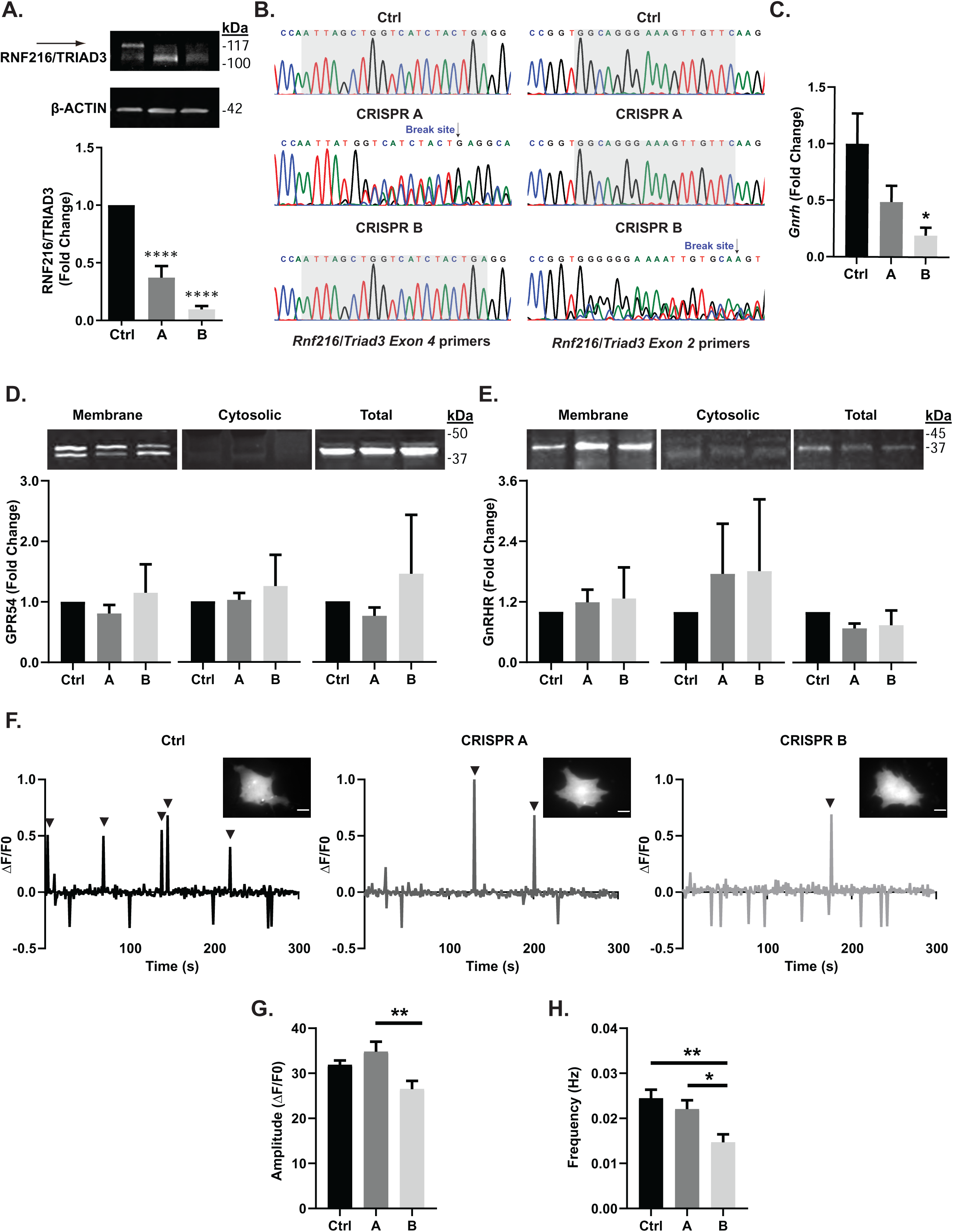
*Rnf216/Triad3* reduces *GnRH* and Ca^2+^ transient frequency in GT1-7 cells. **(A)** Generation of *Rnf216/Triad3* hypothalamic GT1-7 knockout cells. *Top*, Representative immunoblot illustrating RNF216/TRIAD3 in CRISPR-Cas9 control (Ctrl) and knockout cells (A and B). *Bottom*, Mean RNF216/TRIAD3 in control and knockout cells. RNF216/TRIAD3 values were normalized to β-ACTIN. (F (2, 15) = 60.31, *p* <0.0001, One-way ANOVA). Bonferroni’s multiple comparisons test show that CRISPR A and B are significantly lower than control (\*\*\*\**p*<0.0001), *N*=6. **(B)** Sanger sequencing demonstrating successful targeting of *Rnf216/Triad3* in CRISPR A (*left*) and B (*right*) with arrows indicating break sites. Highlighted region represents gRNA targeting site/s. **(C)** Quantitative PCR demonstrating a significant decrease in *Gnrh* (F (2, 6) = 5.129, *p*=0.05, One-way ANOVA). Bonferroni’s multiple comparisons test showed a non-significant reduction in CRISPR A by 51.36% ± 0.1445%, *p*=0.1820 and a significant reduction in CRISPR B by 80.82% ± 0.06998%, \**p*=0.0389. *N*=3. **(D-E)** Subcellular fractionation of control and *Rnf216/Triad3* knockout cells. Representative immunoblots of membrane, cytosol, and total fractions of GPR54 and GnRHR in Ctrl and knockout cells. *N*=4. **(F)** Calcium signaling in Ctrl and *Rnf216/Triad3* knockout cells. Representative fluorescence intensity plots of CRISPR Ctrl, A, and B. Positive signals were measured as 2 standard deviations above the baseline mean indicated by (▾). *Inset,* Representative fluorescent images from each condition. Scale bars represent 10 µm. **(G)** Average amplitude of positive event transients. F (2, 81) = 5.690, *p*=0.0049. One-way ANOVA with Tukey post-hoc analysis. \**p*<0.05, \*\**p*<0.005. **(H)** Frequency of event transients counted as the total number of positive signals in 300 seconds. (F (2, 81) = 7.263, *p*=0.0013, One-way ANOVA) with Tukey post-hoc analysis. *N*=28 for Ctrl, CRISPR A, and CRISPR B. Error bars are represented as + SEM.

Previously, we found that Triad3A, an expressed isoform from the *RNF216/TRIAD3* gene, associates with clathrin-coated pits and alters AMPA receptor endocytosis through ubiquitination of the plasticity associated protein ARC (Mabb et al., 2014; Wall et al., 2018). Modulation of Triad3A also led to global changes in receptor turnover rates in primary hippocampal neurons (data not shown). Thus, we reasoned that the reduction of *Gnrh* could be due to RNF216/TRIAD3-mediated trafficking of key HPG axis receptors or even through ubiquitination of ARC. GT1-7 cells endogenously express GPR54, GnRH receptors (GnRHR), (Mellon et al., 1990; Tonsfeldt et al., 2011) and have also been described to express AMPA receptor subunits (Mahesh et al., 1999; Spergel et al., 1994). However, we were unable to detect membrane expression of the major AMPA receptor subunits, GluA1 and GluA2 in this cell line (data not shown). Therefore, we assessed changes in membrane, cytosolic, and total cell fractions of the key HPG axis receptors GPR54 and GnRHR; but, found no significant differences in the membrane fractions of these receptors when normalized to total protein (Figure 1D-E and Supplementary Figure 1C). We next measured estrogen receptor alpha (ERα), a member of the nuclear receptor family, which has been shown to suppress GnRH production in the presence of estradiol (Otani et al., 2009). However, we did not find any differences in total levels of ERα or the previously identified Triad3A substrate, ARC (Supplementary Figure 1D-E). To determine if the activity of these cells was altered as a result of the loss of *Rnf216/Triad3*, we transfected the genetically encoded calcium indicator, R-GECO1 (Zhao et al., 2011) into each CRISPR cell line (Figure 1F). Although there were no differences in event amplitudes between control and CRISPR B (Figure 1G), there was a significant reduction in event frequency in CRISPR B compared to control (*p*= 0.0013, One-way ANOVA; Figure 1H) and CRISPR A (*p*= 0.0194). These findings suggest that loss of *Rnf216/Triad3* results in decreased *Gnrh* and cell activity. The lack of changes in localization and expression of key HPG axis receptors suggest unknown mechanisms that reduce *<u>Gnrh</u>* and Ca^2+^ transients in GT1-7 cells.

### Generation of *Rnf216/Triad3* KO mice

We next sought to evaluate the role of *Rnf216/Triad3* within an intact HPG axis by generating a constitutive *Rnf216/Triad3* knockout (KO) mouse. *Rnf216/Triad3^fl/fl^* mice were crossed with the female homozygous CMV-Cre global deleter line (*CMV-X^Cre^X^Cre^*) (Schwenk et al., 1995) and the Cre was bred out to avoid phenotypic effects (Supplementary Figure 2A). Genotypes of mice were validated (Supplementary Figure 2B) and corresponding representative immunoblots demonstrated a significant loss of RNF216/TRIAD3 in the hypothalamus and gonads of KO mice (Figure 2A). Although there was a significant decrease in body weight in 4-week old male KO mice, we found no significant difference in body weights in male and female KO mice up to 52 weeks of age (Figure 2B). Normalized brain weights also showed a main effect of sex in 16- (*p*<0.0001, Two-way ANOVA; Figure 2C) and 52-week-old mice (*p*=0.0004), but there were no genotypic differences. Similar to our findings in *Rnf216/Triad3* KO GT1-7 cells (Figure 1D), there were also no differences in expression of GPR54, GnRHR, ERα, or ARC in the hypothalamus of KO mice from both sexes (Supplementary Figure 2C-F).

**Figure 2:**
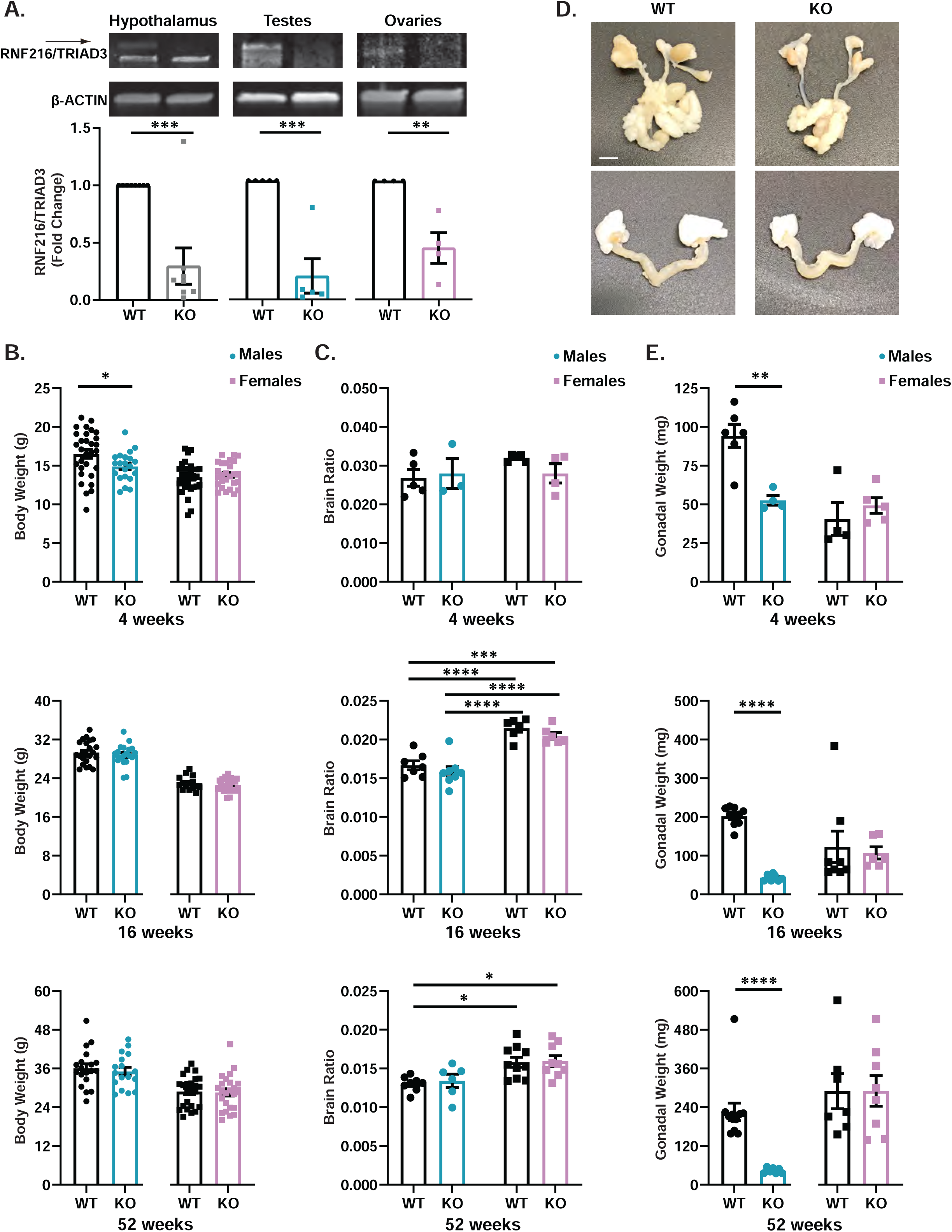
Male gonadal underdevelopment in *Rnf216/Triad3* knockout mice. (A) Representative immunoblots of RNF216/TRIAD3 in WT and *Rnf216/Triad3^-/-^* constitutive knockout (KO) mice. RNF216/TRIAD3 values were normalized to β-ACTIN. KO animals show decreased expression in the hypothalamus (*t* (14) = 4.454, \*\*\**p*=0.0005), testes (*t* (8) = 5.526, \*\*\**p*=0.0006), and ovaries (*t* (6) =4.383, \*\**p*=0.0047); Unpaired *t*-test. *N*= 8 for hypothalamus. *N*= 4 for testes and ovaries. **(B)** Body weights of WT and KO mice at 4-, 16-, and 52-weeks. Male KO mice show decreased body weight at 4-weeks (t (49) = 2.157, \**p*=0.0360); Unpaired t-test. *N*=21-30 per sex/genotype. No significant differences at 16- or 52-weeks was observed. *N*=19-24 per sex/genotype. **(C)** Brain weights of WT and KO mice at 4-, 16-, and 52-weeks. Brain weight was normalized to body weight. There was a main effect of sex at 16- (F (1, 23) = 66.11, *p*<0.0001) and 52-weeks (F (1, 29) = 16.25, *p*=0.0004). Two-way ANOVA with Tukey post-hoc analysis. *N*=3-8 per group. \**p*<0.05, \*\*\**p*<0.0005, \*\*\*\**p*<0.0001. **(D)** Representative images of male and female reproductive organs in WT and KO mice at 16-weeks of age. Scale bar represents 25 mm. **(E)** Gonadal weights of WT and KO mice at 4-, 16-, and 52-weeks. Male KO mice show a significant reduction in testicular weights compared to WT at 4- (*t* (8) =4.321, \*\**p*=0.0025), 16- (*t* (13) =13.96, \*\*\*\**p*<0.0001), and 52-weeks (*t* (17) =5.065, \*\*\*\**p*<0.0001). No significant differences in ovarian weights. Unpaired t-test. *N*=9-12 per group. Error bars are represented as ± SEM.

### *Rnf216/Triad3* KO mice have reproductive impairments

As stated previously, individuals with GHS display HH and reproductive impairments. Upon further inspection of the reproductive organs (Figure 2D), we found that male KO animals had dramatically reduced testicular weights compared to WT at 4- (*p*=0.0025, unpaired-t-test), 16- (*p*<0.0001), and 52-weeks (*p*<0.0001; Figure 2E). However, this was not evident in ovaries isolated from KO females (Figures 2D and 2E). We next determined if the altered testicular weight in males affected breeding viability. We paired WT male:KO female, KO male:WT female, KO male:KO female, and HET male:HET female mice for a maximum of 90 days (Figure 3A). In all pairings, there were no significant differences in the number of days before the first litter, indicating all genotypes had the capacity to produce litters (Figure 3A, left). Nevertheless, there were significant differences in the fertility index, which was calculated as the number of litters produced by each cage (*p*= 0.005, One-way ANOVA; Figure3A, middle). WT male:KO Female pairings had significantly more litters than KO male:WT female (*p*=0.0270) and KO male:KO female (*p*=0.0037) pairings. HET male:HET female pairings had significantly more litters than KO male:WT female (*p*=0.0104) and KO male:KO female (*p*=0.0013) pairings. Notably, there were no differences between HET male:HET female and WT male:KO female pairings. Within litters, we also observed a significant difference in the total number of pups across all litters (*p*= 0.0006, One-way ANOVA; Figure 3A, right). KO male:KO female pairings produced a significantly lower number of pups than WT male:KO female (*p*=0.0006) and HET male:HET female pairings (*p*=0.004). We also found that some litters did not survive past P7; therefore, we also assessed pup survival (Figure 3B). Out of the 77.86% of pups that survived, only 1.56% originating from KO male breeders and 23.44% from KO female breeders successfully grew to weaning age at P21. Surprisingly, of the 22.14% of pups that did not survive, 34.48% of pups came from KO male breeders and the 55.17% of pups came from KO female breeders, suggesting that KO females do not provide adequate maternal care within the first week of postpartum.

**Figure 3:**
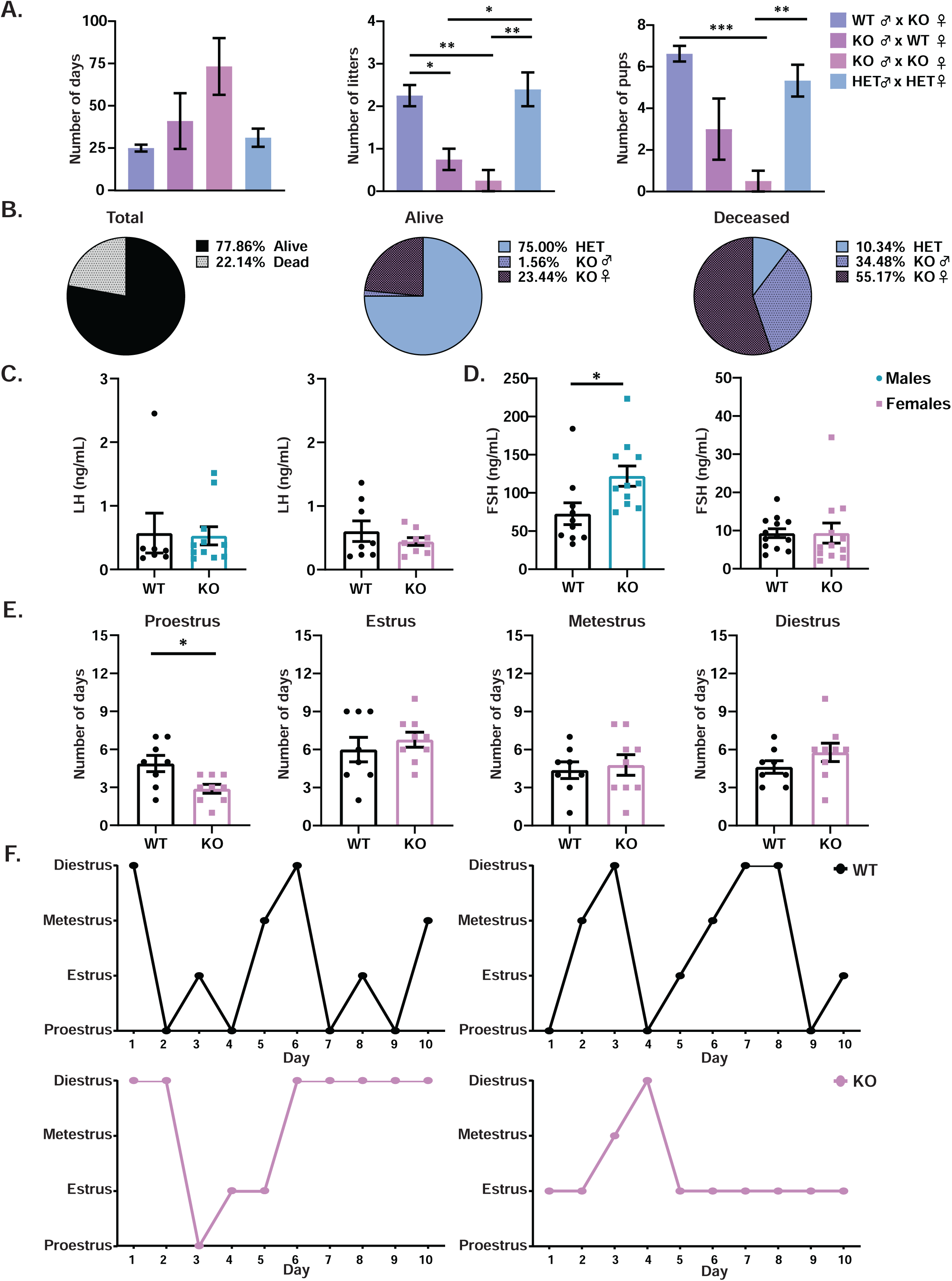
Deletion of *Rnf216/Triad3* induces sex differences in reproduction and FSH release. (A) Assessments of breeding viability in WT and KO mice at 6-7 weeks. *Left*, there were no differences in the number of days before the first litter was produced. There were significant differences in fertility indexes (F (3, 13) =11.76, *p*=0.0005) (*middle*) and number of pups per litter (*right*) (F (3, 24) = 8.213, *p*=0.0006). One-way ANOVA with Tukey’s post hoc analysis. \**p*<0.05, \*\**p*<0.005, \*\*\**p*<0.001, \*\*\*\**p*<0.0001, *N*=4-5 cages per genotype crossing and *N*=8-12 pups per litter. **(B)** Percentage of pup survival from WT and KO breeding pairs. n=102 for pups that survived and n=29 for pups that died. **(C)** No differences in LH serum levels in WT and KO mice at 16-weeks. **(D)** Male KO animals have elevated FSH (*left*) (*t* (19) =2.521, \**p*=0.0208), but females (*right*) show no differences. **(E)** Days females spent in each estrous phase for a duration of 20 days. KO females spent significantly less time in proestrus compared to WT (*t* (15) =2.809, \**p*=0.0132); Unpaired t-test. *N*= 8-9. Error bars are represented as ± SEM. **(F)** Representative estrous cycle of WT (black) and KO (pink) females. KO females showed irregular cyclicity compared to WT females.

Due to these alterations in breeding, we also determined if there was an associated change in HPG axis hormones that could explain the reductions in testicular weight in male KO and decreased maternal care in female KO mice. Blood sera levels of the pituitary hormones FSH and LH were measured in males and females at 16-weeks of age. Surprisingly, neither males nor females exhibited alterations in baseline LH (Figure 3C). While WT and KO females had no changes in FSH, KO males had a significant increase in circulating FSH (*p*=0.0208, unpaired t-test; Figure 3D) compared to WT males, suggesting a causative role for increased FSH in reduced testicular weights and infertility. Although there were no hormonal changes between WT and KO females, when evaluating the estrous cycle for 20 days, KO females spent significantly less time in proestrus (*p*=0.0132, unpaired t-test, Figure 3E). Moreover, WT females cycled regularly throughout the different estrous phases while KO females were arrested in particular phases such as diestrus (Figure 3F).

### *Rnf216/Triad3* KO mice have altered GnRH morphology

In view of *Rnf216/Triad3* KO mice displaying sex differences in reproductive impairment and the variability of GHS patient responses to exogenous GnRH treatment (Margolin et al., 2013; Seminara et al., 2002), we sought to determine if GnRH neurons within the HPG axis were altered *in vivo*. GnRH neurons undergo a migratory and maturation process from the nasal placode to the hypothalamus during embryonic and postnatal development (Cottrell et al., 2006; Jasoni et al., 2009). A recent study using GN11 cells, an immature GnRH neuronal cell line, suggested that migration of GnRH neurons to the hypothalamus could be reduced upon depletion of RNF216/TRIAD3 (Li et al., 2019). Thus, we evaluated if GnRH cell density and morphologies were disrupted within the preoptic area of the hypothalamus in adult KO mice. Histological analysis of the preoptic area of the hypothalamus in male and female KO mice showed no significant differences in the density of GnRH neurons or number of dendrites protruding off the soma (Figure 4A and 4B). We next categorized GnRH neurons by dendritic morphologies based on their number of dendrites which reflects their maturity. For instance, in the hypothalamus, mature GnRH neurons are characterized as having unipolar and bipolar dendritic morphologies whereas immature GnRH neurons contain multiple branches (> 2) (Cottrell et al., 2006). We classified GnRH cells as none (zero dendrites), unipolar (1 dendrite), bipolar (2 dendrites), and multipolar (>2 dendrites) (Supplementary Figure 4) as previously described (Tata et al., 2017). Although KO mice exhibited a higher percentage of none type GnRH neurons (12.78% in males) and (16.07% in females), compared to WT (8.11% in males) and (7.97% in females), there was not a significant difference as assessed by a Chi-squared analysis (Figure 4C). On the contrary, we found a difference in the size of GnRH neurons, which was significantly reduced in KO males (*p*<0.0001, unpaired *t*-test; Figure 4D left) and females (*p*= 0.0002; Figure 4D right). Moreover, there was a significant decrease in the integrated density of GnRH in both KO males (*p*=0.0001, unpaired *t*-test; Figure 4D right) and females (*p*=0.0018) indicating that they may contain less GnRH, which aligned with our *in vitro* data (Figure 1C). Taken together, these findings demonstrate that the loss of RNF216/TRIAD3 initially induces subtle decreases in baseline GnRH in both males and females. We propose a model whereby the loss of RNF216/TRIAD3 results in reduced GnRH neuron activity that causes a modest decrease in GnRH expression in both males and females (Figure 4E). This dysfunction may alter unknown downstream pathways or feedback loops within the HPG axis that lead to sex-specific alterations in reproductive function and behavior.

**Figure 4:**
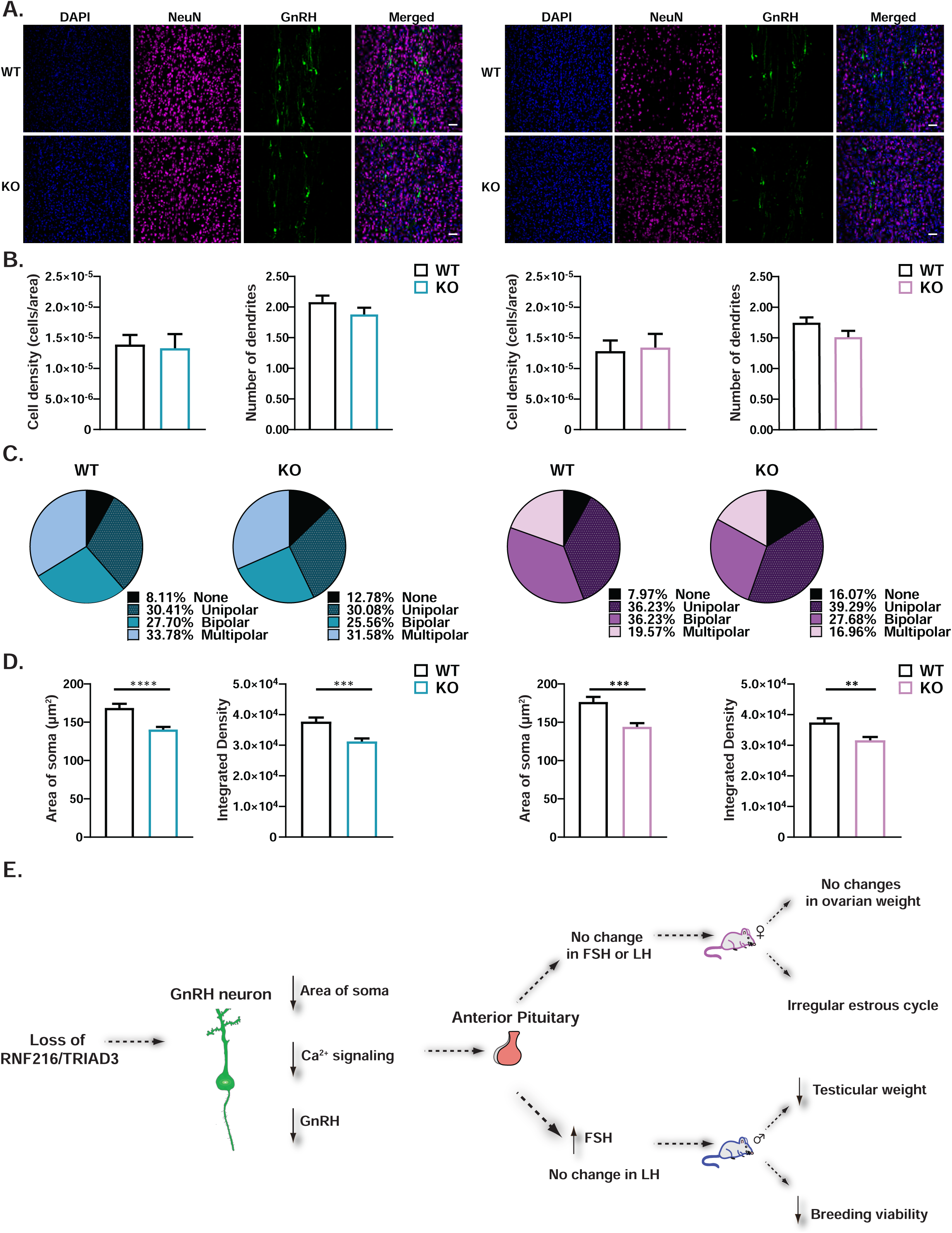
Loss of RNF216/TRIAD3 decreases GnRH soma size and GnRH production in both sexes. (A) Representative confocal images of GnRH cells in the preoptic area of the hypothalamus in adult WT and KO male (*left*) and female (*right*) mice. GnRH neurons were imaged at 20X magnification. Scale bars represent 50 µm. **(B)** No differences in the number of dendrites and cell density in both sexes. **(C)** GnRH neurons were classified according to the number of dendrites protruding directly off the soma: none (zero dendrites), unipolar (1 dendrite), bipolar (2 dendrites), multipolar (>2 dendrites). In both sexes, KO animals had a higher percentage of none type compared to WT. This was not significantly different in males (X^2^(3) = 1.709, *p*=0.6349) or females (X^2^(3) = 5.198, *p*=0.1579). Chi-squared. **(D)** Significant differences in soma area in males (*t* (281) =4.182, \*\*\*\**p*<0.0001) and females (*t* (248) =3.821, \*\*\**p*=0.0002) compared to respective WT counterparts. There were also significant differences in the integrated density in males (*t* (281) =3.853, \*\*\*\**p*=0.0001) and females (*t* (248) =3.162, \*\**p*=0.0018); Unpaired t-test. *N*=3 for males, 149 WT and 134 KO cells across 3 sections per animal. *N*=3 for females, 138 WT and 112 KO cells across 3 sections per animal. Error bars are represented as + SEM. **(E)** Model depicting the loss of RNF216/TRIAD3 on HPG axis function. Depletion of RNF216/TRIAD3 leads to decreased GnRH neuron volume, *GnRH* production, and activity. These GnRH changes could result in sex differences in gonadotropin release. In males, alterations lead to reduced testicular weights and decreased breeding viability and females display irregular estrous cycles without changes in ovarian weights.

## Discussion

RNF216/TRIAD3 is a multifunctional E3 ligase that ubiquitinates numerous substrates involved in inflammation and immunity, (Chuang and Ulevitch, 2004; Fearns et al., 2006; Nakhaei et al., 2009), autophagy (Kim et al., 2018; Wang et al., 2016; Xu et al., 2014), and synaptic plasticity (Mabb et al., 2014; Wall et al., 2018). Despite the divergent roles of RNF216/TRIAD3, its position within the neuroendocrine system has not been elucidated. Here, we found that the loss of RNF216/TRIAD3 results in reproductive impairment at multiple levels within the HPG axis, affecting males and females differently. Abnormal GnRH and GnRH neuron sizes were identified in both males and females; yet, only KO males showed changes in FSH, gonadal development, and breeding viability. On the other hand, KO females did not show these phenotypes, but did have modest estrous cycle abnormalities.

Due to the role of TRIAD3A in the central nervous system, we sought to determine if loss of RNF216/TRIAD3 affected GnRH neuron activity. Using the CRISPR-Cas9 system in a GnRH hypothalamic-derived cell line (GT1-7), we generated two *Rnf216/Triad3* KO cell lines. One cell line (CRISPR B) resulted in a complete knockout of *Rnf216/Triad3* whereas the other (CRISPR A) still retained a small fraction of WT sequence (Figures 1A-B, Supplementary Figure 1A-B). *Gnrh* was significantly reduced in CRISPR B (Figure 1C) which coincided with reduced baseline calcium frequency (Figure 1G). The cell autonomous feature of RNF216/TRIAD3 parallels our *in vivo* data illustrating abnormal GnRH expression and decreased soma size (Figure 4). These findings indicate that complete removal of RNF216/TRIAD3 is required for achieving a full phenotypic effect and is consistent with the recessive nature of inheritance for GHS (Alqwaifly and Bohlega, 2016; Margolin et al., 2013).

RNF216/TRIAD3 disease-causing mutations decrease its E3 enzymatic activity and disrupt ARC ubiquitination, providing strong evidence that ubiquitination of substrates is a major contributing factor for GHS (Husain et al., 2017; Mabb et al., 2014). However, we did not find evidence of changes in ARC in GT1-7 KO cells or the hypothalamus of *Rnf216/Triad3* KO mice (Figure 1 and Supplementary Figure 1D-E, Supplementary Figure 2). Moreover, a transgenic mouse mutation of RNF216/TRIAD3 Arc ubiquitination sites did not cause reproductive impairments in male and female mice (Wall et al., 2018), providing strong evidence for RNF216/TRIAD3-dependent regulation of other substrates that regulate HPG axis function. Triad3A can ubiquitinate Arc, which stimulates AMPA receptor trafficking in the hippocampus (Mabb et al., 2014), and knockdown of Triad3A decreased receptor trafficking (data not shown). Therefore, we determined if RNF216/TRIAD3 can alter membrane localization of GPR54, a kisspeptin receptor found in GnRH cells that precedes the release of GnRH. However, we found no differences in membrane localization of GPR54 and GnRHR in *Rnf216/Triad3* KO cells (Figure 1D and Supplementary Figures 1D-E)) nor did we observe changes in ERα, and ARC at steady state (Figure 1 and Supplementary Figure 1D-E) and *in vivo* (Supplementary Figure 2). Given our inability to identify targets, then, what could be potential RNF216/TRIAD3 substrates in GnRH neurons?

One possible substrate is Beclin-1, which is ubiquitinated by RNF216/TRIAD3 to mediate autophagy (Kim et al., 2018; Wang et al., 2016; Xu et al., 2014). RNAi-mediated silencing of *Rnf216/Triad3* increased Beclin-1 and inhibited GnRH cell migration, which was found to function independently of ARC (Li et al., 2019). Our results demonstrate that there are no changes in GnRH neuron density in both WT and KO adult mice of both sexes. KO mice also expressed varying types of GnRH morphologies that included mature and immature GnRH neurons in the preoptic area of the hypothalamus (Figure 4C, Supplementary Figure 4), demonstrating that there were no gross impairments on GnRH neuron migration and dendritic morphologies. The loss of *Becn1* in the perinatal ovary leads to loss of female germ cells and infertility (Gawriluk et al., 2011). However, Beclin-1 has never been linked to GnRH neuron morphology and cell activity or GnRH production, suggesting that there are other possible substrates that are ubiquitinated by RNF216/TRIAD3 to regulate GnRH neuron functions.

There are many factors involved in the signaling and production of GnRH, one of which is nuclear factor kappa-light-chain-enhancer of activated B cells (NF-κB)-dependent signaling (Zhang et al., 2013). RNF216/TRIAD3 has been shown to ubiquitinate multiple proteins within the innate immunity pathway, which alter NF-κB activity (Alturki et al., 2018; Chuang and Ulevitch, 2004; Fearns et al., 2006; Nakhaei et al., 2009). Interestingly, both NF-κB and IKK-β activation reduce GnRH release from GT1-7 cells and decrease *Gnrh1 in vivo*, which is reversed upon NF-κB inhibition (Zhang et al., 2013). Thus, it is possible that RNF216/TRIAD3 ubiquitinates factors within the NF-κB pathway to positively regulate GnRH release. Although we did not find changes in GPR54 expression, the altered calcium signaling we observed suggests that there are other intrinsic factors that RNF216/TRIAD3 interacts with to affect baseline calcium responses. Depolarization of GnRH neurons activates the phospholipase C-β signaling pathway that engages transient receptor potential channels (TRPC) (Rønnekleiv and Kelly, 2013; Zhang et al., 2008). GnRH transcription and secretion is increased through hydrolysis of phosphatidylinositol 4,5- bisphosphate (PIP_2_), which leads to Ca^2+^ mobilization and activation of protein kinase C (PKC)(Wetsel et al., 1993; Wetsel and Negro-Vilar, 1989). GnRH release is also affected by voltage-gated Ca^2+^ channels that include L-type and T-type channels that modulate burst firing in GnRH neurons through estradiol stimulation (Lee et al., 2010; Watanabe et al., 2004; Zhang et al., 2009). Ventral anterior homeobox1 (VAX1) is a homeodomain protein that also controls *Gnrh1* transcription (Hoffmann et al., 2016). Deletion of *Vax1* leads to delayed puberty, hypogonadism, and infertility. The effects of VAX1 on GnRH production are known to depend on other transcriptions factors such as SIX6. It is therefore possible that RNF216/TRIAD3 ubiquitinates these factors to regulate *GnRH*.

As we moved downstream within the HPG axis, we found that deletion of *Rnf216/Triad3* led to sex differences in gonadotropin hormone release and reproductive function. KO males exhibited decreased testicular weight throughout their lifetime (Figure 2D-E) and reduced breeding viability (Figure 3A) similar to findings in a different *Rnf216* mouse model (Melnick et al., 2019). Although KO females did not show a reduction in ovarian weights, they displayed irregular estrous cyclicity and spent less time in proestrus (Figure 3F-G), the phase essential for copulation. Sex differences in reproductive function were also apparent in GHS patients with different point mutations within *RNF216/TRIAD3*. For example, male patients with mutations in the c.2061G>A location demonstrate poor development of secondary sexual characteristics, hypogenitalism, gynecomastia, and low LH and testosterone with no changes in FSH (Alqwaifly and Bohlega, 2016). Similarly, male patients with mutations within the catalytic region (R751C) also display no detectable LH secretion, while males with compound heterozygous mutations in the E205fsX15 + C597X catalytic region demonstrate orchidopexy at 6 years old and low testosterone concentration in adulthood (Margolin et al., 2013; Mehmood et al., 2017). A female patient with the R751C mutation had menarche at 16 years old followed by secondary amenorrhea with no detectable LH secretion whereas another female patient with a p.G138GfsX74 mutation, outside the catalytic domain, had oligomenorrhea followed by amenorrhea at 27 years old with low-amplitude LH secretion (Margolin et al., 2013). Surprisingly, we did not observe changes in circulating LH secretion at one time point, but this could be due to the pulsatility of LH secretion as these clinical studies measured endogenous LH at an hourly rate. It is likely that individual point mutations within regions of the *RNF216/TRIAD3* gene create broad phenotypes of hypogonadotropic hypogonadism while also displaying nuances in reproductive impairment.

KO male mice displayed increased FSH (Figure 3D) without changes in LH (Figure 3C), whereas females did not show differences in release of pituitary hormones. Because no sex differences in depolarization of GnRH neurons have been reported (Liu et al., 2008), it is likely that sex differences in gonadotropin hormone release occurs either at the level of GnRH release or within gonadotropes. Notably, GnRH regulates the release of FSH and LH from gonadotropes via distinct mechanisms (Kile and Nett, 1994). It is also possible that male gonadotropes are more sensitive to subtle changes in GnRH. The sex differences in FSH release and subsequent reproductive function could also point to sexual dimorphism in gonadotropes from the anterior pituitary. There is only one study that provides evidence for dimorphism in gonadotropes. Although expressed in both male and female gonadotropes, FSH selectively colocalizes with neuronal Ca^2+^ sensor synaptotagmin-9 (syt-9) in females to facilitate FSH exocytosis. Deletion of *syt-9* in mice decreases basal and stimulated FSH secretion exclusively in females and alters their estrous cycle, without any effect on males (Roper et al., 2015). It is likely that there are additional sex-specific molecular factors that regulate FSH release in males, for which RNF216/TRIAD3 may be a likely candidate.

FSH is known to have different functions in males and females indicating that the sex-specific reproductive phenotypes in *Rnf216/Triad3* KO mice could be due to factors involved in gonadal feedback loops. FSH is critical for spermatogenesis in males but stimulates the ovarian follicle in females. Inhibin A and B are glycoproteins that control the synthesis and release of FSH secreted in a gonadal-dependent way (Groome et al., 1996; Kubini et al., 2000; Luisi et al., 2005; Welt et al., 1997) and clinically, men with HH display low levels of inhibin B (Coutant et al., 2010). Subsequent testosterone and estradiol release from the gonads also provides negative feedback to regions of the hypothalamus and pituitary (Welt et al., 2005). Cumulative imbalances of inhibin A and B, testosterone, and estradiol could also explain the sex differences seen in *Rnf216/Triad3* KO mice.

## Conclusion

Our findings suggest that effects of RNF216/TRIAD3 loss occur at different points within the HPG axis. Future experiments would be tailored to evaluating changes in isolated gonadotropes and HPG axis feedback loop hormones in KO mice. One way to reconcile RNF216/TRIAD3-specific actions would be to spatially control the removal of RNF216/TRIAD3 within select nodes of the HPG axis. Our work highlights the importance of ubiquitination in proper HPG axis function. By establishing areas of impairment and the substrates involved, it may be possible to identify sex-specific targets for GHS and other neuroendocrine disorders related to ubiquitin disruption.

## Acknowledgements

We would like to acknowledge Zachary Allen, for help in development, genotyping, and maintenance of the *Rnf216/Triad3* KO mice; Antoinette Charles, for cloning assistance and validation of gRNA targets; and Dr. Wei Wei, for training in immunohistological techniques. We would also like to thank Georgia State University Neuroscience Institute department members, Dr. Daniel Cox for use of the LightCycler instrument; Claudia Sanabria for training in confocal techniques in the Confocal Core Facility; and Drs. Nancy Forger and Alexandra Castillo-Ruiz for experimental design consultation and feedback. This work was funded by a National Ataxia Foundation Young Investigator Research Grant, the Cleon C. Arrington Research Initiation Grant Program (RIG-93), and a Molecular Basis of Disease Seed Grant to A.M.M.; the National Science Foundation of Hungary (K128317) and the Hungarian Brain Research Program (2017-1.2.1-NKP-2017-00002) to E.H.; National Institutes of Health grant (R01GM115763) and Georgia State University startup funds to N.F.; Y.C.H. and A.J.G. were funded by a Brains & Behavior Fellowship; A.J.G. was also funded by the Kenneth W. and Georganne F. Honeycutt Fellowship.

## Author Contributions

A.J.G. conceived and performed all the experiments including generation and maintenance of GT1-7 CRISPR clones and *Rnf216/Triad3* mice, conducted the data acquisition and analysis, and wrote and edited the manuscript. Y.C.H. and C.W. cloned the *Rnf216/Triad3* CRISPR plasmids and edited the manuscript. B.D. and N.F. provided instrumentation, performed the calcium imaging, and edited the manuscript. E.H. provided the GnRH antibody, assisted with experimental design, and edited the manuscript. A.M.M. conceived the experiments, generated the *Rnf216/Triad3* mouse colony, collected blood samples for hormonal analysis and brains for histological analysis, wrote and edited the manuscript, and supervised the study.

## Methods

### Animals

Mice were kept in standard housing with littermates, provided with food and water ad libitum and maintained on a 12:12 (light-dark) cycle. All behavioral tests were conducted in accordance with the National Institutes of Health Guidelines for the Use of Animals. Mice were treated in accordance with the Animal Welfare and Ethics Committee (AWERB) and experiments were performed under the appropriated project licenses with local and national ethical approval. Samples sizes for behavior and immunohistochemistry experiments were calculated using variance from previous experiments to indicate power, which statistical analysis for significance was set at 95%. All behavioral studies and isolation of body tissue for biochemical experiments, vaginal cytology, and blood collection were approved by the Georgia State University Institutional Animal Care and Use Committee.

### Generation of Rnf216/Triad3 knockout mice

Embryonic stem cell clones were generated to target exons 3 to 4 of the *Rnf216/Triad3* gene on mouse chromosome 5, which prevents production of all isoforms (International Knockout Mouse Consortium). ES cell clones were injected into blastocytes and implanted into pseudopregnant mice. These mice were crossed to heterozygous mice for Flp recombinase to excise out the LacZ/neomycin cassette in order to obtain one *Rnf216* allele flanked by loxP (fl) sites to generate C57BL/6N-Rnf216<tm1c(EUCOMM)Wtsi>/Tcp (*Rnf216/Triad3^wt/fl^*) (Canadian Mouse Mutant Repository at the Hospital for Sick Children, Toronto, CA). Homozygous floxed conditional male mice (*Rnf216/Triad3^fl/fl^*) were crossed with homozygous CMV-Cre^+/+^ female mice (The Jackson Laboratory, JAX stock #006054) to allow the cre to excise out exons 3 to 4, creating a dysfunctional gene. To breed out the cre, female offspring that were heterozygous for loxP and cre (*Rnf216/Triad3^wt/fl^::*CMV-cre^-/+^) were bred with WT males to generate heterozygous *Rnf216/Triad3*^-/+^ mice. Male *Rnf216/Triad3*^-/+^ mice were then bred with WT females to generate *Rnf216/Triad3*^-/+^ mice. *Rnf216/Triad3*^-/+^ male and female mice were then bred together to generate the experimental animals used for this study (*Rnf216/Triad3*^+/+^ (WT), *Rnf216/Triad3*^-/+^ (HET), and *Rnf216/Triad3*^-/-^ (KO)).

### Blood collection and ELISA

Male bedding was inserted in female cages 48 hours prior to blood collection to synchronize estrous cycles and vaginal cytology was performed prior to blood collection. Blood samples were collected (BD Microtainer Gold, ThermoFisher) at 11:00am during each session through retro-orbital bleeds, tail vein, or after decapitation and stored at room temperature for 30-60 minutes. Samples were then centrifuged at room temperature at 3,500 rpm for 10 minutes. Serum was collected and stored at -80°C until further study. Serum samples were shipped to the University of Virginia Center for Research in Reproduction, Ligand Assay and Analysis Core (supported by the Eunice Kennedy Shriver NICHD Grant R24 HD102061) for LH and FSH analysis.

### Vaginal cytology

Female mice at age 16 weeks underwent vaginal lavage with sterile Dulbecco’s phosphate-buffered saline (DPBS) at 11:00am for 20 consecutive days. Cells were collected, mounted on coverslides, and stained with thionin (0.01 g/mL thionin acetate, 0.2% acetic acid, 0.007 g/mL sodium acetate anhydrous, pH 4.25) for 10 minutes and washed briefly with ddH_2_O. and examined under light microscopy to determine phase of the cycle using the classification protocol described in (Byers et al., 2012).

### Breeding viability

The following pairings were set up to assess breeding viability: WT male with KO female; KO male with WT female; KO male with KO female; and HET male with HET female. Animals were between 6-8 weeks old prior to pairing. Females were primed with male bedding 48 hrs prior to pairing. Breeding cages contained 1 male and 1 female for a duration of 90 days. The number of litters per cage, the number of pups per litter per cage, and the number of days before the first litter were measured. The litters and pups were counted between P0-P1. These assessments also included litters and pups that did not survive through weaning at P21.

### Immunohistochemistry

Mice were perfused with room temperature 4% paraformaldehyde/1X PBS. Brains were collected, post-fixed in 4% PFA/1XPBS overnight at 4°C, and then placed in sucrose sinking solution (30% sucrose in 1X PBS) for 24hrs. Brains were embedded with embedding matrix (M-1 Embedding Matrix, Thermo Fisher) and then frozen at -80°C. Embedded brains were equilibrated at -20°C for at least 1 hour and then sectioned at 30 µm thickness using a cryostat (Lieca CM 3050S) and stored in freezing solution (45% PBS, 30% ethylene glycol, 25% glycerol) at -20°C. Stereologically matched sections of the preoptic area, medial preoptic area, anteroventral periventricular nucleus, and arcuate brain sections were selected and washed 3x at room temperature in 1XPBS and blocked with blocking buffer (10% normal goat serum and 0.5% Triton X-100) overnight at 4°C. The following primary antibodies were added in blocking buffer and incubated for 24 hours at 4°C: 1:500 dilution of Rat polyclonal anti-GnRH antibody (Skrapits et al., 2015) and 1:250 dilution Guinea Pig polycloncal anti-NeuN (EMD Millipore). After the brain slices were washed 3x with 1XPBS they were incubated with the following secondary antibodies in blocking buffer for 2 hours at room temperature: 1:500 dilution of Donkey anti-rat Alexa Fluor 488 (Jackson ImmunoResearch Laboratories), 1:200 dilution of Donkey anti-guinea pig Alexa Fluor 647 (VWR), and 1:500 dilution of DAPI (4’,6-Diamidino-2-Phenylindole, Dihydrochloride) (ThermoFisher). The brain slices were washed 3x with 1XPBS and then mounted with mounting media (Fluoro-Gel, VWR) on coverslips (Superfrost™ Plus Microscope Slides, ThermoFisher) for imaging. Fixed slides were imaged on a laser scanning confocal microscope (Zeiss, LSM 700) using a 20x Plan-Apochromat N.A. 0.8 without oil immersion and 40x Plan-Apochromat N.A. 1.4 with oil immersion objectives. GnRH was excited using a 488 nm laser, NeuN was excited using a 639 nm laser, and DAPI was excited using a 405 nm laser. Z-stack projections were obtained at 1µm intervals (11 slices) and images were displayed as maximum intensity projections of the entire z-series using Zen black software (Zeiss). After image collection, GnRH neurons were analyzed as described previously in (Cottrell et al., 2006) for cell soma area, total number of dendrites projecting from the soma, number of dendritic branch points, defined as the number of points of dendritic bifurcation giving rise to an additional process length of more than 5 μm. GnRH neuronal morphologies were classified as described previously in (Tata et al., 2017) as mature or immature using the following criteria: unipolar/mature (one dendrite directly off the GnRH soma), bipolar/mature (two dendrites directly off the GnRH soma), or complex/immature (three or more dendritic processes directly off of the GnRH soma). Values of quantified GnRH dendritic morphologies were expressed as the percentage of the total GnRH neuron population analyzed.

### Cloning of control and Rnf216/Triad3 CRISPRs

The type II CRISPR nuclease system was implemented in GT1-7 cells to facilitate genome editing by co-expressing a codon-optimized cas9 nuclease along with a single guide RNA (sgRNA). The LentiCRISPR (pXPR_001) plasmid contains two expression cassettes, hSpCas9, and the chimeric guide RNA (Shalem et al., 2014). Briefly, 5µg of lentiviral CRISPR plasmid was digested with *BsmBI* (NEB) for 90 min at 37°C. After electrophoresis of the digested vector, the 11 kB band was gel purified using the QIAquick Gel Extraction Kit (QIAGEN) according to the manufacturer’s protocol. Two sgRNA sequences targeting mouse *Rnf216/Triad3* were designed using http://crispr.mit.edu: CRISPR A (TCAGTAGATGACCAGCTAAT) which targets exon 4 and CRISPR B (GAACAACTTTCCCTGCCACC) which targets exon 2. Primer sequences are as follows:

CRISPR A-F 5’-CACCGTCAGTAGATGACCAGCTAAT-3’;

CRISPR A-R 5’-AAACATTAGCTGGTCATCTACTGAC-3’;

CRISPR B-F 5’-CACCGGAACAACTTTCCCTGCCACC-3’;

CRISPR B-R 5’-AAACGGTGGCAGGGAAAGTTGTTCC-3’.

DNA oligos were phosphorylated and annealed by mixing each oligo pair (100µM) with 10X T4 Ligation Buffer (NEB), ddH_2_O, and T4 PNK (NEB). The reaction underwent the following cycling conditions: 37°C for 30 min, 95°C for 5 min, and then ramped down to 25°C at 0.1°C/sec. Products were then diluted 1:200 in elution buffer and finally cloned into the vector using a ligation reaction that included the *BsmBI* digested plasmid, diluted oligo, 2X Quick Ligase Buffer (NEB), ddH_2_O, and Quick Ligase (NEB) that was incubated at room temperature for 10 min. The targeting guide sequences were transformed into competent cells and positive clones were sequenced for insert validation. Knockout efficiency for each individual targeting sequence was validated by transient transfection and immunoblotting in the B16 mouse melanoma cell line. CRISPR clones A and B were used to generate *Rnf216/Triad3* knockout clones in GT1-7 cells.

### Cell Culture and generation of control and Rnf216/Triad3 CRISPR clones

GT1-7 cells were a kind gift from Dr. Pamela Mellon (Mellon et al., 1990). Cells were maintained in DMEM (Corning) with 10% FBS (GE Healthcare) and 1% Pen Strep (ThermoFisher) at 37°C with 5% CO_2_. Cells were seeded at a density of 2.5 x 10^5^ cells/well in a 6-well dish and transfected with CRISPR-Cas9 plasmids using the standard Lipofectamine 3000 (Thermo Fisher) protocol. Briefly, 2.5µg of plasmid DNA was mixed with Lipofectamine 3000 reagents and incubated for 24 hours before replacing with fresh media. After 48 hours, 2.0 µg/mL of puromycin was added to the media for selection. After integration was established, single cell clones were selected and expanded into colonies under puromycin selection. Loss of *Rnf216/Triad3* was validated through immunoblotting. CRISPR clones that showed a loss of RNF216/TRIAD3 were used for subsequent experiments.

### Validation of CRISPR sequences

Confirming targeted CRISPR sequences were adapted from the Giuliano et al. protocol (Giuliano et al., 2019). GT1-7 CRISPR cells were plated at a density of 1.0×10^6^ in a 6-well dish. Genomic DNA was extracted using the DNeasy Blood & Tissue Kit (QIAGEN) according to manufacturer’s protocol. The concentration and purity of purified DNA was assessed using a NanoDrop ND-2000/2000c (ThermoFisher). DNA was only used if it had an A260/A280 ratio between 1.8–2.0. PCR of CRISPR control, A, and B was performed using specific primer sets for CRISPR A and B. PCR products were run on a 2% agarose gel and gel bands were manually extracted and purified using QIAquick PCR & Gel Cleanup Kit (QIAGEN) according to manufacturer’s protocol. Purified PCR products were submitted to GENEWIZ (South Plainfield, NJ) for Sanger sequencing. Sequencing results were analyzed using the previously established Tracking of Indels by DEcomposition (TIDE) software (https://tide.nki.nl/) (Brinkman et al., 2014). Control CRISPR sequences were used as reference chromatograms for CRISPR A and B comparison. Primer sequences for gRNA targeted *Rnf216/Triad3* genomic regions are as follows:

CRISPR A-F 5’-ATGGCGGAAAAACATTGGGC-3’;

CRISPR A-R 5’-ACCTGGACCAAGCAGTAAGC-3’;

CRISPR B-F 5’-AACAGTAGAATCGCTCTGGCT-3’;

CRISPR B-R 5’-CTTGTTTTTCAAACCCTGCAGAAC-3’

### RT-qPCR

GT1-7 cells were seeded at a density of 2.5×10^5^ in a 6-well dish. QIAzol (QIAGEN) reagent from the RNeasy Lipid Tissue Mini Kit (QIAGEN) was directly added onto the plates, removed using a cell scraper (Corning), and passed through a 27G needle. The concentration and purity of purified RNA was assessed spectrophotometrically using the NanoDrop ND-2000/2000c (ThermoFisher). RNA was only used if it had an A260/A280 ratio between 1.8–2.25. First-strand cDNA synthesis was performed using the iScript Reverse Transcription Supermix for RT-qPCR (BioRad) according to the manufacturer’s protocol. RT-qPCR was performed using the Real-Time PCR System (LightCycler 96, Roche). Each reaction comprised of 0.5 µL of diluted cDNA, 5 µL FastStart Essential DNA Green Master Mix (Roche) and 10 µM primers in a final volume of 10

µL. The PCR cycling conditions were as follows: activation at 95°C for 900s; then 3-step amplification with 45 cycles of 95°C for 15s; 63°C for 15s; and 72°C for 60s. Cycling was followed by melt curve recording at 95°C for 10s; 65°C for 60s; and 97°C for 1s. Primer standard curves were performed to estimate the PCR efficiencies for each primer pair. Cycle threshold (Ct) values were determined by Lightcycler 96 application software. All qPCR reactions were run in triplicate with at least 3 biological replicates. A mean Ct value was calculated for each primer pair and each experimental condition. Relative quantification of *GnRH* mRNA was performed using the 2-ΔΔCt method (Livak and Schmittgen, 2001). Data were normalized to the geometric mean of *Gadph* and presented as expression relative to a standard condition as indicated in the figure legends. *GnRH* and *Gapdh* primers were designed using sequences from previous studies. Primer sequences are as follows:

GnRH-F: 5’-GCTCCAGCCAGCACTGGTCCTA-3’;

GnRH-R: 5’-TGATCCACCTCCTTGCCCATCTCTT-3’ (Nuruddin et al., 2014);

GAPDH-F: 5’-GGCAAATTCAACGGCACAGT-3’;

GAPDH-R: 5’-GGGTCTCGCTCCTGGAAGAT-3’ (Wall et al., 2018).

### Subcellular fractionation

Fractionation of GT1-7 cells were adapted from Abcam (https://www.abcam.com/protocols/subcellular-fractionation-protocol). Each cell line was seeded at a density of 1×10^7^ cells on a 10 cm dish. After the media was removed, ice cold DPBS was added to the plates to wash off remaining media and dead cells. 500µL of fractionation buffer (Abcam) was added and the cells were removed by scraping. Membrane, nuclear, cytoplasmic, and mitochondrial fractions were isolated according to the manufacturer’s protocol. Cell fractions were stored in -80°C until processed. Cell fraction samples were thawed on ice and 10-20 µL of RIPA buffer (50mM Tris-HCl pH 8.0, 150mM NaCl, 1% NP-40, 0.5% sodium deoxycholate, 0.1% SDS) was added for 30 minutes with disrupting the pellet every 5 minutes and then centrifuged for 20 minutes at 14,000 rpm at 4°C. The soluble fraction was collected and 2XSDS sample buffer (10% 1M Tris-HCl pH6.8, 3% DTT, 4% SDS, 20% glycerol, 0.2% Bromophenol blue, 1:1000 dilution of β-Mercaptoethanol) was added to each protein sample and boiled at 95°C for 7 min before loading on an SDS-PAGE gel (see below). After protein transfer, membranes were blotted with Revert 700 total protein stain (LiCor) before blocking.

### Western Blotting

Mouse tissue was thawed on ice and 300-500 µL of RIPA buffer was added and then the tissue was homogenized using sterile pestles. The samples were then centrifuged at 14,000 rpm for 15 minutes at 4°C and the supernatant was collected for protein quantification. GT1-7 cells were prepared as stated previously in the subcellular fractionation section. The soluble fraction was collected and the protein concentration was determined using Pierce 660nM Protein Assay Kit (ThermoFisher). The samples underwent SDS-polyacrylamide gel electrophoresis and were transferred on nitrocellulose membrane (BioRad) for 1hr at 70 mV. The blots were incubated overnight at 4°C with blocking buffer (Intercept (TBS) blocking buffer, LiCor). Membranes were then probed with the following primary antibodies prepared in a 1:1 ratio of TBST (1X TBS, 0.1% Tween-20) and blocking buffer solution with 1:1,000 20% NaN_3_: rabbit polyclonal anti-RNF216 (Bethyl Laboratories, 1:1,000), rabbit anti-kiss1/GPR54 (Lifespan Biosciences, 1:600), rabbit anti-GnRHR (Lifespan Biosciences, 1:600), rabbit anti-Arc (Synaptic Systems, 1:1,000), mouse anti-β-Actin (Genetex, 1:3,000) and were incubated overnight at 4°C. The following secondary antibody dilutions were prepared in a 1:1 ratio of TBST (1X TBS, 0.1% Tween-20) and blocking buffer solution with 1:2,000 20% SDS: IRDye 680RD Goat anti-Mouse IgG (H+L) (Li-COR, 1:20,000) and IRDye 800CW Goat anti-Rabbit IgG (H+L) (Li-COR, 1:15,000) and were incubated for 1hr at room temperature. Blots were imaged using the Odyssey CLx Imaging System (LI-COR) with a resolution of 169 µm, medium quality and a 0 mm focus offset. Images were processed using the Gel Analysis tool in Image J using individual channels. Briefly, boxes were drawn around each band. Once the lanes were labeled and plotted, the area of the peaks were selected and measured. For each blot, proteins of interest were normalized to a loading control.

### Calcium Imaging

GT1–7 CRISPR cells were plated on glass cover-slips (22 mm diameter) coated with poly-D-lysine (100 µg/mL) at a density of 30,000 cells/coverslip. The cells were maintained as previously described under the cell culture section. The next day, cells were transfected with CMV-R_GECO (Addgene #45494) (Wu et al., 2013) using Lipofectamine 3000 as described above. During imaging, cells were shifted to ACSF solution containing 118 NaCl, 3 KCl, 0.5 CaCl_2_, 6 MgCl_2_, 5 HEPES, 25 NaHCO_3_, 11 D-glucose, pH 7.3. Cells were imaged in live conditions using highly inclined and laminated optical sheet (HILO) microscopy on a Nikon Ti-E inverted microscope equipped with a Nikon 100× TIRF objective and an Andor IXon Ultra 897 EMCCD camera at an imaging speed of 1 frame per second. A CW 561 nm laser (Oxxius) was used for exciting the fluorophore and a Quad Band filter set (TRF89901v2, Chroma) was used for rejecting the fluorescence background. Calcium signals were acquired after measuring ΔF/F using the “delta F up function” and the “F div F0” function in Image J (FIJI). Values were normalized to one using (x – min(x)) / (max(x) – min(x)). Peak amplitudes were extracted using findpeaks in Matlab (The MathWorks, Inc.), which were defined as 2 standard deviations above the mean fluorescence.

### Statistical Analysis

Statistical analyses applied were the post hoc Student’s t test, One-way ANOVA, and Two-way ANOVA with multiple comparisons. Bonferonni and Tukey’s test were used for comparing group means only when a significant F value was determined. For all comparisons, significance was set at p < 0.05. Data presented in figures and tables are means (± SEM).

---

**Figure S1:**
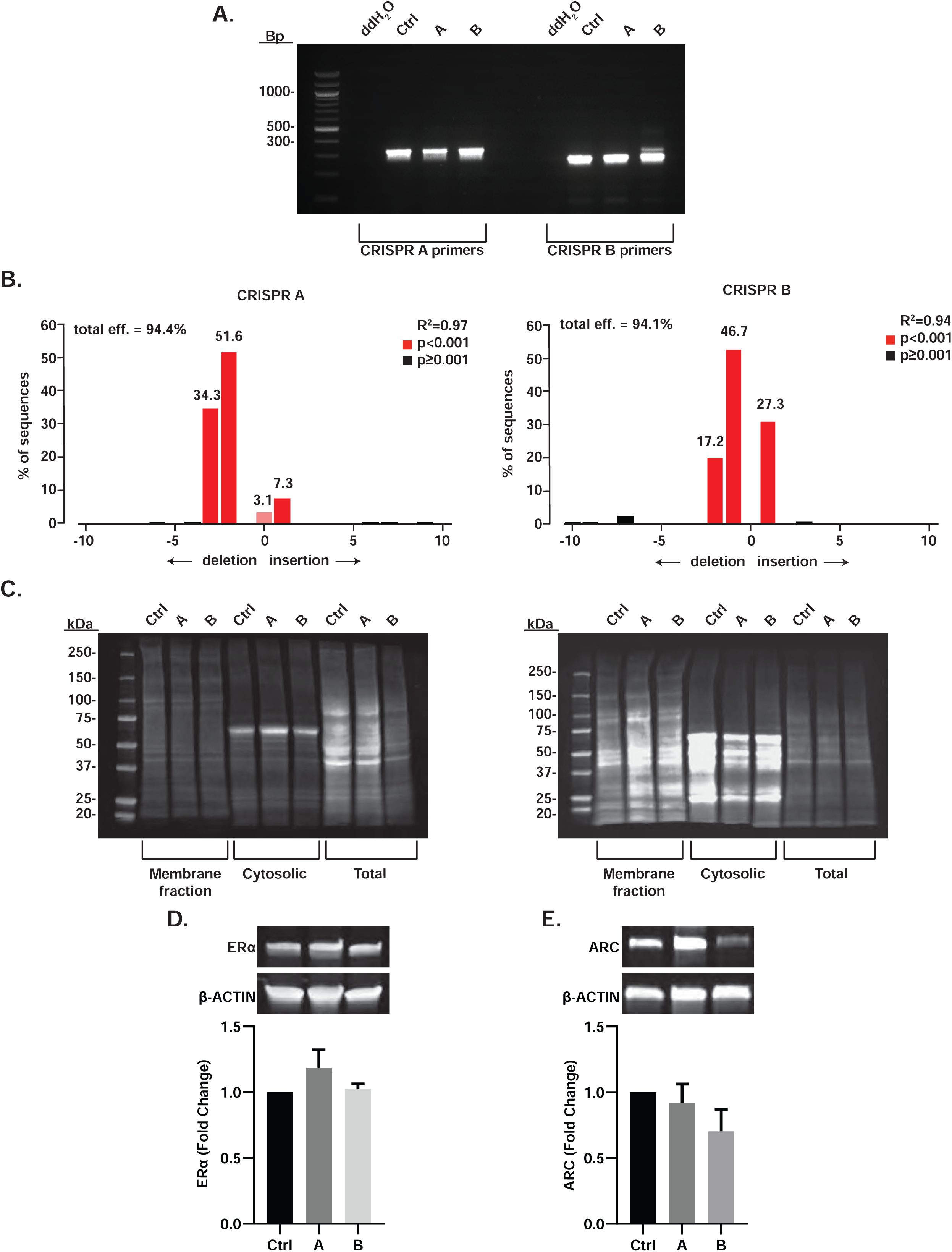
**(A)** PCR analysis of GT1-7 CRISPR control (Ctrl), A, and B using specific primer sets for CRISPR A and B. CRISPR A primers produced a product length of ∼330bp, and CRISPR B primers at ∼260bp. **(B)** Identification of induced mutations in CRISPR A and CRISPR B using TIDE analysis software. In this non-linear model, R^2^ is the goodness of fit, and the p-value threshold is set at *p*=0.001 and is associated with the estimated abundance of each indel, calculated by a two-tailed *t*-test of the variance–covariance matrix of the standard errors. **(C)** Total protein stain from Ctrl and *Rnf216/Triad3* knockout cells in membrane, cytosolic, and total fractions for GPR54 (*left*) and GnRHR (*right*). **(D)** Representative immunoblots illustrating ERα in Ctrl and knockout cells (A and B). *Bottom*, Respective protein values were normalized to β-ACTIN. There was no significance between groups. N = 3 for each group. **(E)** Representative immunoblots illustrating ARC in CRISPR-Cas9 Ctrl and knockout cells (A and B). *Bottom*, Respective protein values were normalized to β-ACTIN. There was no significance between groups. N = 6 for each group. Error bars are represented as + SEM.

**Figure S2:**
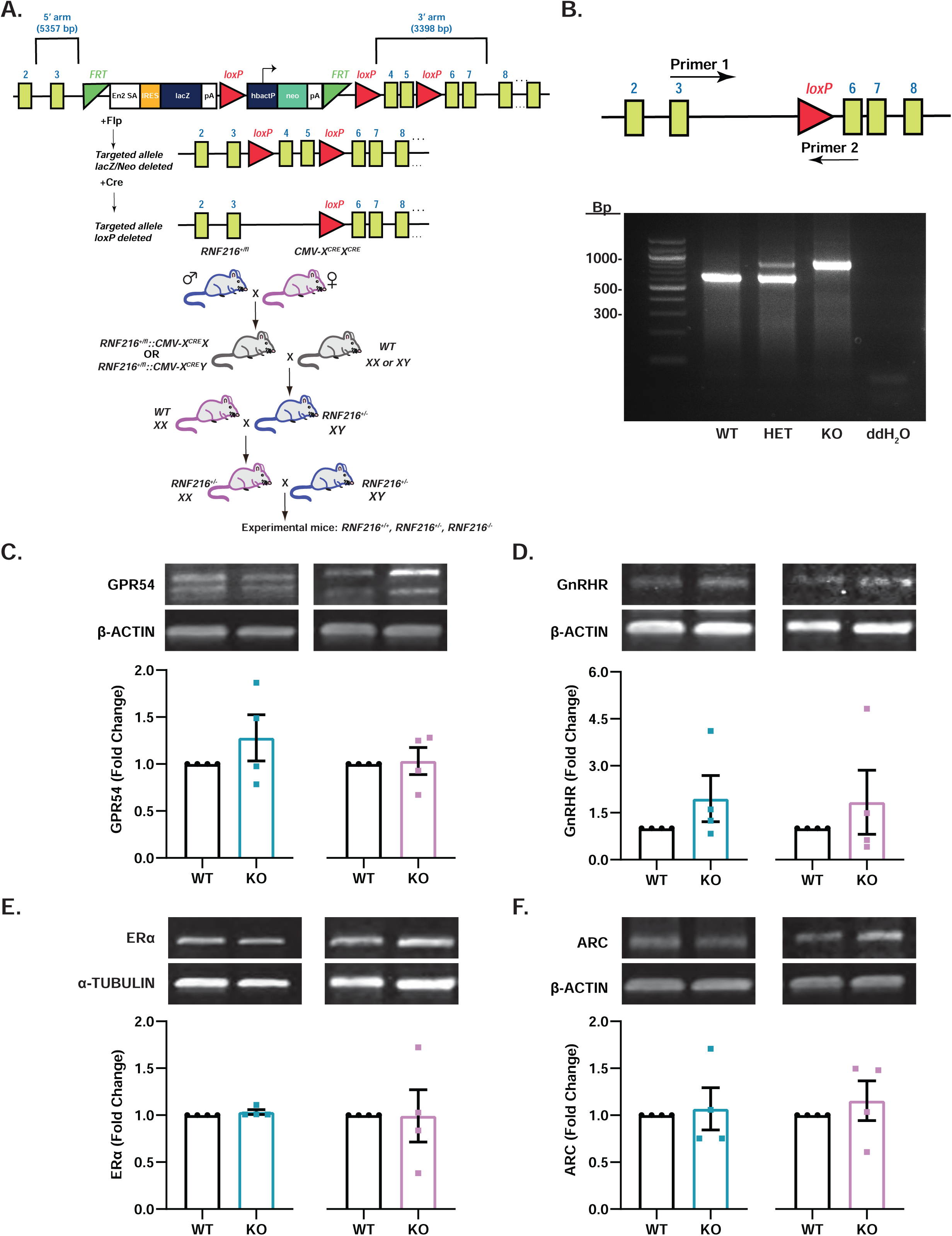
**(A)** Generation of *Rnf216/Triad3* constitutive knockout (KO) mice. *Rnf216^fl/fl^* mice were crossed with the female homozygous CMV-Cre global deleter line (*CMV-X^Cre^X^Cre^*). Male offspring that were heterozygous for *Rnf216^fl/+^* were bred with wildtype (WT) females. *Rnf216^-/+^* male and female offspring were then bred together to generate wildtype (*Rnf216/Triad3^+/+^*), heterozygous *(Rnf216/Triad3^-/+^*), and knockout (*Rnf216/Triad3^-/-^)* mice. **(B)** Genetic validation of *Rnf216/Triad3* KO mice. *Left*, Primer positions represent sites used to distinguish wildtype (WT), heterozygous (HET), and homozygous (KO) mice. *Bottom*, Genotyping results from WT, HET, and KO mice. Wildtype band size is 624 base pairs and the floxed band size is 890 base pairs. **(C-F)** Protein expression in hypothalamus of *Rnf216/Triad3* mice for GPR54, GnRHR, ERα, and ARC. Respective protein values were normalized to either β-ACTIN or α-TUBULIN. There were no differences between WT and KO. N=8 per genotype. Error bars are represented as ± SEM.

**Figure S3:**
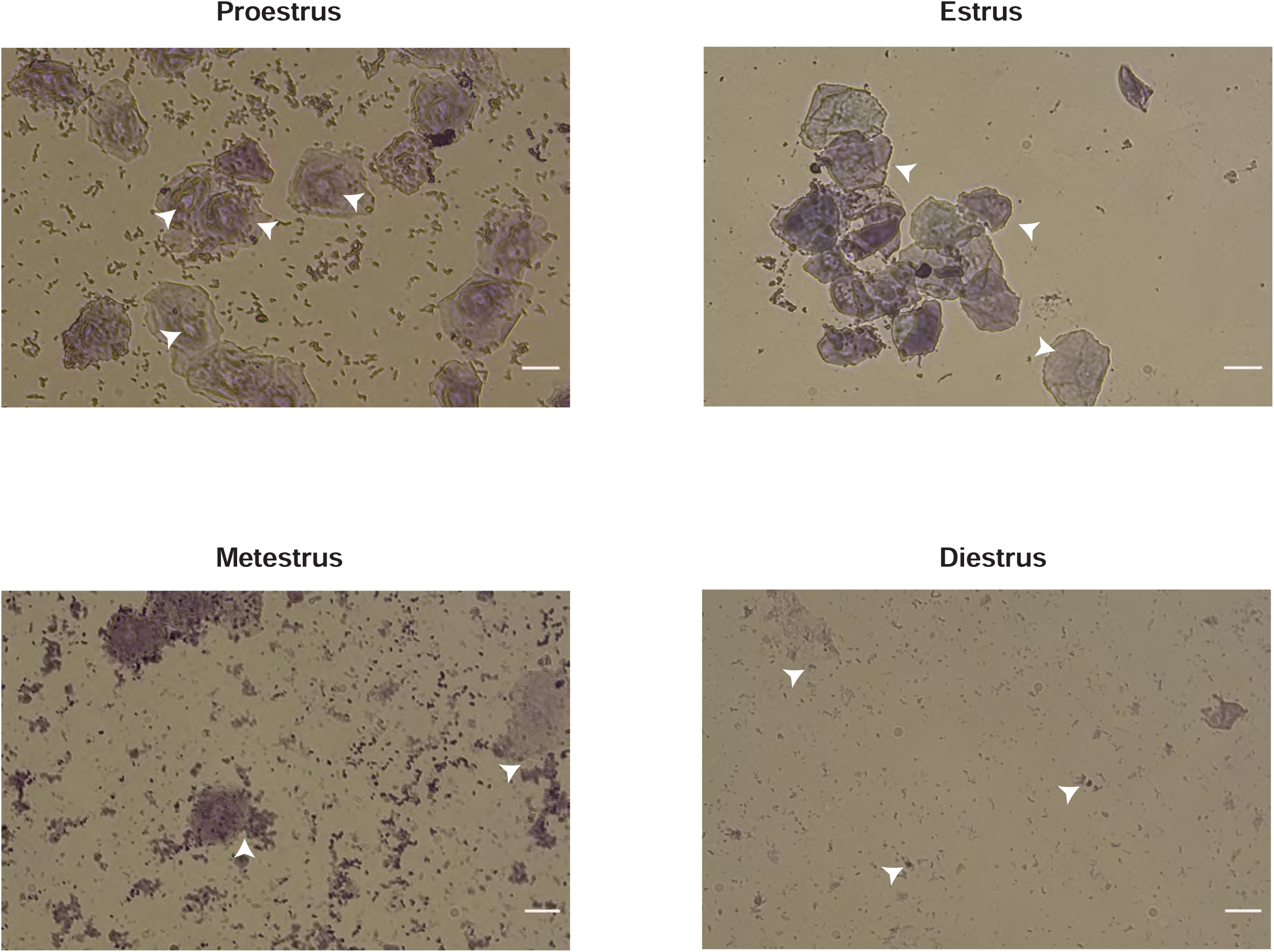
Representative images of each phase of the estrous cycle. Cells were visualized with a thionin stain and classified according to the specific cell type/s for each phase indicated by white arrows. Proestrus is identified through nucleated epithelial cells. Estrus contain cornified epithelial cells. Metestrus show both cornified epithelial cells with leukocytes. Diestrus consists of a majority of leukocytes. Scale bars represent 100 µm.

**Figure S4:**
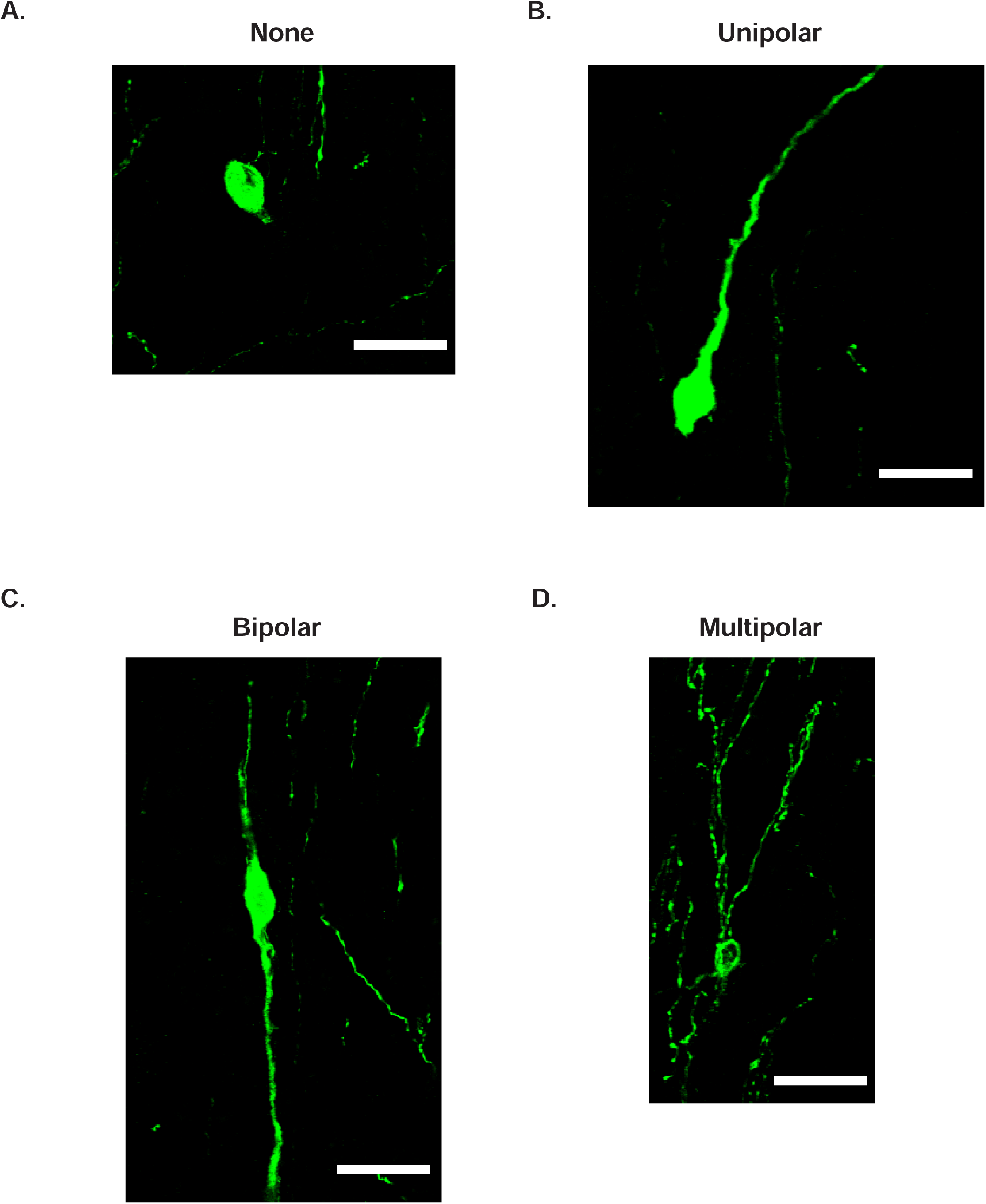
Representative confocal images of each identified GnRH morphology type in the preoptic area of the hypothalamus of WT animals. **(A)** None, GnRH neurons that had no dendrites. **(A)** Unipolar, GnRH neurons that had 1 dendrite protruding off the soma. **(C)** Bipolar, GnRH neurons that had 2 dendrites protruding off the soma. **(D)** Multipolar, GnRH neurons that had >2 dendrites protruding off the soma. Scale bars represent 25 µm.

